# DEAR-O: Differential Expression Analysis based on RNA-seq data - Online

**DOI:** 10.1101/069807

**Authors:** Zong-Hong Zhang, Naomi R. Wray, Qiong-Yi Zhao

## Abstract

**Summary:** Differential expression analysis using high-throughput RNA sequencing (RNA-seq) data is widely applied in transcriptomic studies and many software tools have been developed for this purpose. Active development of existing popular tools, together with emergence of new tools means that studies comparing the performance of differential expression analysis methods become rapidly out-of-date. In order to enable researchers to evaluate new and updated software in a timely manner, we developed DEAR-O, a user-friendly platform for performance evaluation of differential expression analysis based on RNA-seq data. The platform currently includes four of the most popular tools: DESeq, DESeq2, edgeR and Cuffdiff2. Based on the DEAR-O platform, researchers can evaluate the performance of different tools, or the same tool with different versions, with a customised number of biological replicates using already curated RNA-seq datasets. We also initiated an online forum for discussion of RNA-seq differential expression analysis. Through this forum, new useful tools and benchmarking datasets can be introduced. Our platform will be actively maintained to ensure new major versions of existing tools and new popular tools are included. DEAR-O will serve the community by providing timely evaluations of tools, versions and number of replicates for RNA-seq differential expression analysis.

**Availability and implementation:** The DEAR-O platform is available at http://cnsgenomics.com/software/dear-o; the online discussion forum is https://groups.google.com/d/forum/dear-o

## 1 Introduction

In RNA-seq experiments, the primary interest of biologists in many studies is differential expression analysis between experimental groups. There are three major steps involved in such analysis: (1) alignment of RNA-seq reads to the reference, (2) estimation of the relative expression level of genes/transcripts by calculating the number of aligned reads (i.e. read counts), and (3) application of statistical methods to test if an observed difference in read counts between two experimental groups is statistically significant (Zhang, et al., 2014). The Negative Binomial (NB) distribution has achieved a dominant position in the methodologies of RNA-seq differential expression analysis as it models the variance in biological replicates very well. A number of popular tools have been developed for differential expression analysis based on the NB distribution (e.g. DESeq (Anders and Huber, 2010), edgeR (Robinson, et al., 2010), DEseq2 (Love, et al., 2014)). Although studies (Rapaport, et al., 2013; Soneson and Delorenzi, 2013; Zhang, et al., 2014) have been conducted to compare the performance of popular differential expression analysis tools and provided useful guidance on the choice of analysis tools, they quickly become out-dated when new versions or new tools are released. For example, since edgeR (Robinson, et al., 2010), Cuffdiff (Trapnell, et al., 2010) and DESeq (Anders and Huber, 2010) were released in 2010, each of these tools have undergone continual upgrades with more than 10 versions. In addition, new algorithms, such as Cuffdiff2 (Trapnell, et al., 2013) and DESeq2 (Love, et al., 2014), which were developed in 2013-14, have both undergone more than 5 upgrades.

The field of RNA-seq differential expression analysis is not short of analysis tools, but there is a lack of timely evaluations and comparisons of tools to deliver useful guidance to end users. To address this issue, we developed DEAR-O, a platform that can serve the RNA-seq community by providing timely evaluations of popular tools and their versions for RNA-seq differential expression analysis. DEAR-O not only includes the comparative analyses we have previously reported (Zhang et al, 2014), but also includes updated comparisons based on upgraded versions of DESeq (4 new versions), edgeR (4 new versions) and Cuffdiff2 (2 new versions) and new algorithm DESeq2 (3 versions), which have been released since our study was published. Another feature of the DEAR-O platform is that researchers can perform evaluations by testing a different number of biological replicates, a key factor in experimental design of RNA-seq studies, for the query and subject sets, separately.

## 2 Functionality and Methods

DEAR-O: Differential Expression Analysis based on RNA-seq data – Online, has been developed in the Perl programming language and HTML. The user input options include choice of data source, choice of analysis tools and their versions, and choice of the number of biological replicates used in the analysis (Supplementary Figure 1). All input fields are pull down selection lists.

Currently, two well-curated publicly available RNA-seq datasets are loaded into DEAR-O: mouse neurosphere data which includes four biological replicates in two treatment groups (Zhang, et al., 2014) and human lymphoblastoid cell lines (LCL) data which includes 40 biological samples from the same population (Pickrell, et al., 2010). For the mouse data, publicly available microarray data for the same treatment groups is used as a gold standard benchmark for gene expression. For the human LCL dataset, we expect no DEGs between two hypothetical groups (i.e. 20 biological samples were randomly selected as one group and the remaining 20 samples formed the other group). We introduced DEGs into real RNA-seq data by simulation as previously described (Zhang et al, 2014). Therefore all DEGs and non-DEGs in the human LCL dataset are known and can be used as a gold standard benchmark.

Major versions of four popular software tools, Cuffdiff2, DESeq, DESeq2 and edgeR are included in the tool list with the release of DEAR-O. After choosing the data source, query tool, subject tool, versions and number of biological replicates, DEAR-O users can click the “Submit” button to start a comparative analysis between the query set (i.e. the query tool, version, number of biological replicates) and the subject set. The detailed process for detecting differentially expressed genes (DEGs) has been described in our previous study (Zhang, et al., 2014). Briefly, TopHat2 (version 2.0.8) (Kim, et al., 2013) was used to align the RNA-seq reads against a reference genome (hg19 for human and mm10 for mouse). Cuffdiff2 was then applied to detect DEGs between experimental groups. HTseq-count (version 0.5p2) was used to generate count tables for DESeq, DESeq2 and edgeR analyses. Based on count tables, DESeq, DESeq2 and edgeR were applied to detect DEGs. The differential expression analysis results were then benchmarked with the gold standard.

To evaluate the performance between the query and subject sets, true positives (TP), false positives (FP), true negatives (TN) and false negatives (FN) are calculated based on the comparison results between RNA-seq data and the gold standard benchmark data. The false positive rate (FPR) and the true positive rate (TPR) are defined as follows: FPR = FP/(FP+TN), TPR = TP/(TP+FN). The receiver operating characteristic (ROC) displays the relationship between TPR and FPR. The area under the ROC curve (AUC) is used to evaluate the performance of the query and subject sets for the detection of DEGs. Two kinds of AUC are included as previously described (Wan and Sun, 2012; Zhang, et al., 2014): (i) AUC1 is the area under the ROC curve in the full range of FPR, i.e. 0≤FPR≤1 and the maximum AUC1 is 1; and (ii) AUC2 is the area under the ROC curve in the range of 0≤FPR≤0.05 and the maximum AUC2 is 0.05.

The output results from the DEAR-O platform include (i) a table to summarise the specificity and sensitivity for the query and subject sets (Supplementary Table 1); (ii) a ROC plot with two AUC values to show the comparative results between the query and subject sets (Supplementary Figure 2); (iii) a table to show the number of DEGs from the query set, the subject set, and the gold standard set (Supplementary Table 2); and (iv) a Venn diagram to show the intersections of the number of DEGs identified by the query, subject and gold standard sets (Supplementary Figure 3).

DEAR-O platform will be actively maintained to include new major versions of existing tools or new popular tools as they become available. We also initiated an online forum for discussion of RNA-seq differential expression analysis. Through this forum, researchers are able to not only have close interactions but also introduce new useful tools and benchmarking datasets to improve the DEAR-O platform.

